# Bead Enhancement of EV Analysis

**DOI:** 10.1101/269423

**Authors:** Hsing-Ying Lin, Katherine S. Yang, Caleigh Curley, Hakho Lee, Marisa W. Welch, Brian M. Wolpin, Ralph Weissleder, Hyungsoon Im, Cesar Castro

## Abstract

Extracellular vesicles (EVs) are recognized cancer biomarkers, however, clinical analysis has been difficult due to a lack of simple and sensitive assays. Here, we describe a bead-enhanced flow cytometry method, BEAD flow, using biotinylated EVs captured on streptavidin particles. With this method, we show analysis of patient-derived EVs using a panel of pancreatic cancer biomarkers. BEAD flow is easily translatable to any biomarker or cancer type and can be run with conventional flow cytometers, making it highly flexible and adaptable to diverse research and clinical needs.

---

Given the advanced and incurable disease often found with pancreatic ductal adenocarcinoma (PDAC) at initial presentation, timely and simple diagnostic methods are needed.^[1]^ Current diagnostic methods are typically invasive and expensive,^[2,3]^ relying heavily on the use of imaging modalities such as CT, MRI, and PET, which often miss early disease. Serum CA19-9 levels are clinically used as diagnostic and predictive biomarkers of PDAC,^[4,5]^ although they often lead to false negative or positive results,^[6,7]^. Methods for noninvasive and early PDAC diagnoses use circulating tumor DNA and cells or pancreatic fluid to identify rare epigenetic changes.^[8]^ Recent work has examined extracellular vesicles (EVs) as potentially invaluable biomarker sources within bodily fluids.^[8–10]^

EVs are a heterogenous population of particles, such as exosomes, microvesicles, and membrane particles that are continuously shed by all cell types, including cancer cells, into circulation.^[11]^ EVs contain protein and RNA cargo highly reflective of their cell of origin, making them invaluable sources of potential biomarkers for liquid biopsy diagnostics.^[12–16]^ Despite the ready accessibility of EVs within bodily fluids, currently no EV analysis methods exist that are sensitive, yet straightforward enough to implement across diverse laboratory settings. Current methods for EV protein analysis rely on insensitive assays such as ELISA and Western blot that require large sample amounts for measurement. Extremely sensitive methodologies are under development by our group and others,^[10,14–19]^ but they often require specialized setups that are not yet commercialized for widespread use.

In response, we sought to develop a standardized method to effectively analyze EV proteins from human samples. To address this unmet need, we developed BEAD (Bead-conjugated EV Assay Detected) flow cytometry, leveraging biotinylated EVs captured on streptavidin-coated polystyrene (PS) beads. Sensitive single EV flow methods are under development, but currently require specialized and dedicated flow cytometers that are not widely available.^[20–22]^ Moreover, labeling and analyzing EVs with antibodies for single particle analysis is expensive, leads to extensive sample loss, and is time consuming. BEAD flow offers several advantages over existing methods i) improved EV capture efficiency due to the high affinity biotin-streptavidin interaction; ii) a simplified assay measured using conventional flow cytometers; iii) enhanced detection sensitivity due to EV (and biomarker) concentration on PS beads. Using BEAD flow, we were able to interrogate pancreatic cancer EV biomarkers in clinical patient samples.

In BEAD flow, EVs are first biotinylated using NHS-PEG4-Biotin (30 min), followed by capture on 5 µm streptavidin-coated polystyrene (PS) particles in another 30 min reaction. Bead-bound EVs are then stained with primary antibodies followed by AlexaFluor 488 secondary antibodies for flow cytometry measurements (**Figure 1**). We used a CytoFlex flow cytometer equipped with automatic handling of samples in a 96-well plate. The entire sample to readout process took approximately 4 hrs for up to 48 samples.

**Figure 1.**
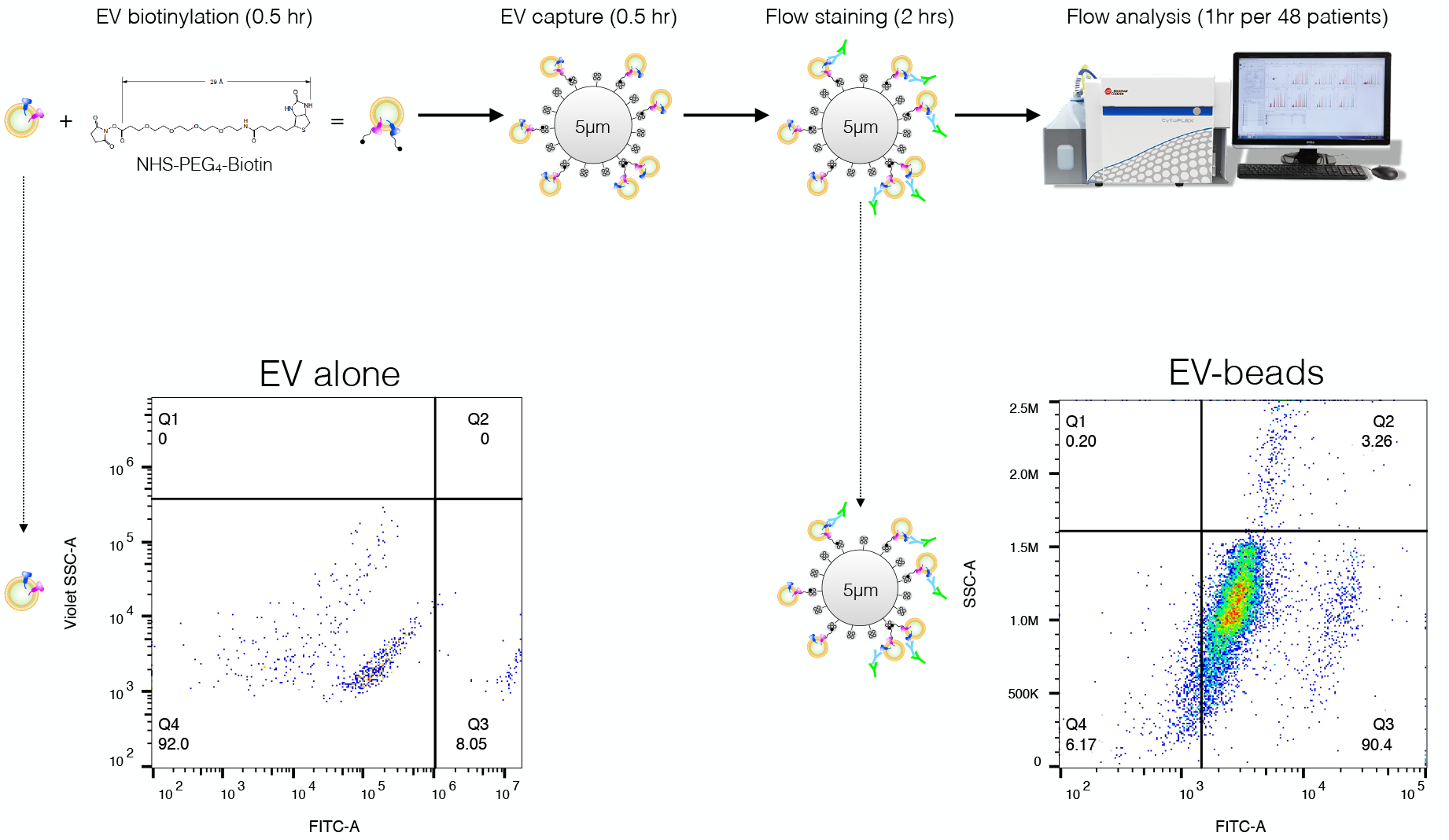
BEAD flow method for patient sample EV analysis. Top) Following isolation from plasma, EVs are biotinylated using NHS-PEG4-Biotin.EVs are captured on 5 µm streptavidin-polystyrene beads and then stained with either primary and AlexaFluor 488 conjugated secondary antibodies or fluorescent primary antibodies. Samples are analyzed on a CytoFlex (Beckman Coulter) 96-well plate flow cytometer. The entire workflow is complete within 4 hrs. Bottom Left) Dot plot of EVs from a patient-derived xenograft cell line (1617 PDAC)stained with a FITC-EGFR antibody. Bottom right) 1617 EVs biotinylated and captured on 5 µm streptavidin polystyrene beads, followed by staining with the same EGFR-FITC antibody as in the bottom left. All dot plots are gated on isotype control stained samples.

Nano-sized EVs exhibit rapid Brownian motion with relatively weak fluorescent signal. Thus, analysis of individual EVs requires a specialized setup and instrument capable of tracking fast-moving EVs with high sensitivity. When we labeled individual EVs isolated from a patient-derived xenograft cell line (1617 PDAC), exhibiting high EGFR expression,^[10]^ with EGFR-FITC antibodies, only 8% of EVs were detected (**Figure 1**). In contrast, when the same EVs were captured on 5 µm PS beads and stained with the same EGFR-FITC antibody, a large signal amplification was observed, resulting in ~90% of EV-bead conjugates positive for EGFR (**Figure 1**).

Previous bead-based EV assays utilized affinity ligands (e.g. antibodies) for marker-based EV capture or passive adsorption of EVs on aldehyde/sulfate latex beads with a hydrophobic surface.^[23,24]^ Capturing biotinylated EVs obviates the need to identify a set of antibodies for capture and labeling and showed improved capture efficiency. The passive EV adsorption on latex beads often requires a minimum 2.5 hr incubation, thus doubling the sample preparation time compared to the strategy described here. To compare the capture efficiencies of latex and streptavidin-coated PS beads, we captured EVs from the same 1617 PDAC cell line on latex or streptavidin-coated PS beads and labeled them with EGFR antibodies. EVs captured on latex beads resulted in only ~7% of beads positive for EGFR, while ~96% of beads were positive for EGFR when biotinylated EVs were captured on streptavidin PS beads (**Figure 2A**). The signal increase with the BEAD flow method was observed with other antibodies against EpCAM and MUC1, as well as a cocktail of antibodies (EpCAM, EGFR, MUC1, WNT-2, GPC1) (**Figure 2B**). Comparing 5 μm aldehyde/sulfate latex beads to same-sized streptavidin PS beads still resulted in slightly better signal for the biotinylated EVs with streptavidin-coated PS beads, likely due to better EV capture on beads through the high affinity biotin-streptavidin interaction (**Figure 2B**). BEAD flow has a detection limit of EV proteins of 74.24 ng, equivalent to 2.29 × 10^7^ EVs (**Figures 2C**, **S1 and** Table S1). BEAD flow also showed good reproducibility with a variation of ~6% in independently repeated experiments (Figures 2D **and S2**).

**Figure 2.**
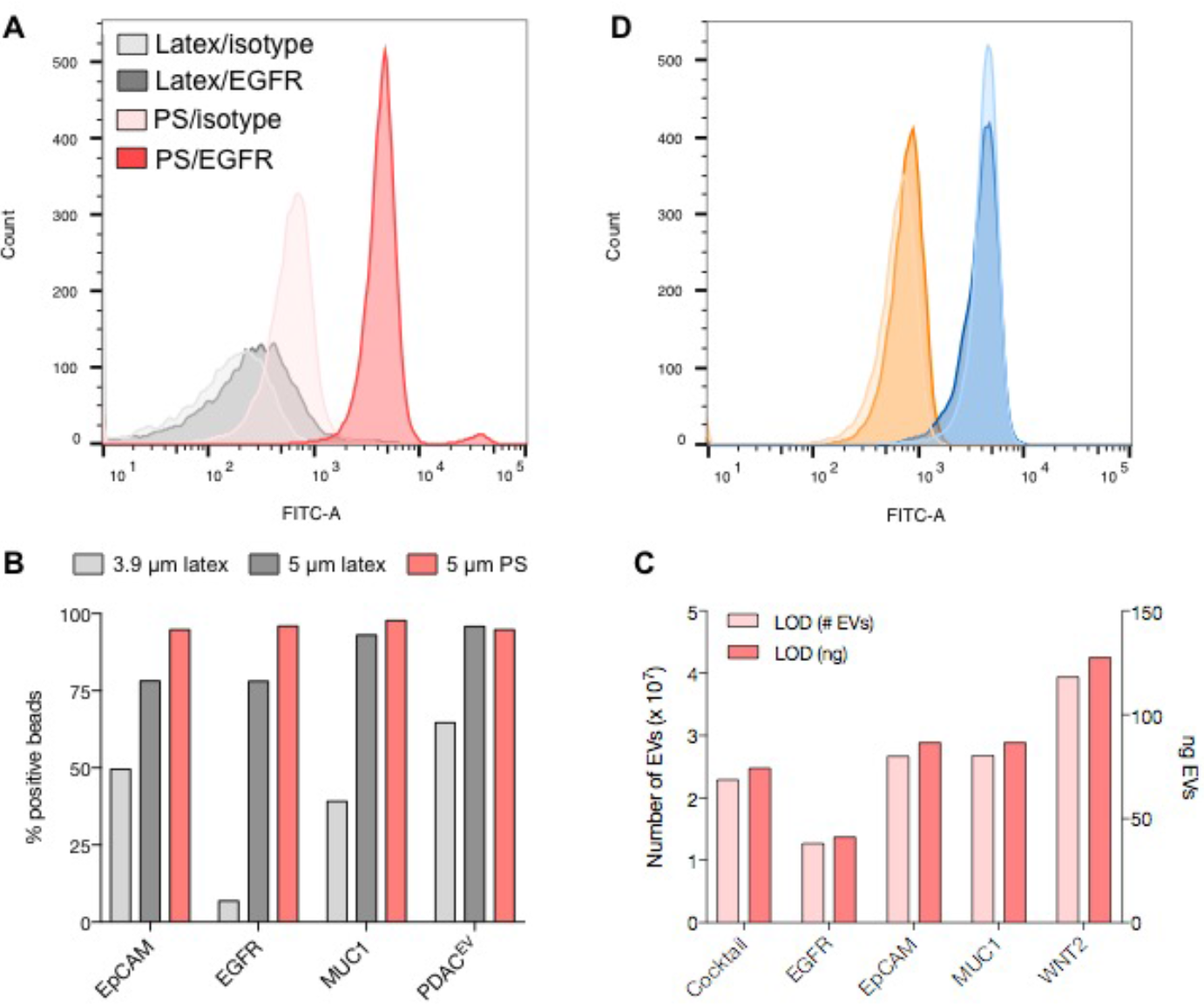
Validation of the BEAD flow method. A) EVs from 1617 PDAC cells were adsorbed onto 3.9 µm aldehyde/sulfate latex beads (gray) or biotinylated and captured on 5 µm streptavidin polystyrene beads (pink). EV-bead conjugates were then stained using an EGFR antibody and AlexaFluor 488 secondary antibody, followed by flow cytometry analysis. B) 1617 PDAC EVs were adsorbed onto 3.9 µm (light gray), 5 µm (dark gray) aldehyde/sulfate latex beads, or biotinylated and captured on 5 µm streptavidin polystyrene beads (pink). EV-bead conjugates were stained with antibodies against EpCAM, EGFR, MUC1, or a five antibody cocktail (PDAC^EV^: EpCAM, EGFR, MUC1, WNT-2, GPC1). Signal from each antibody was gated to an identical EV-bead conjugate stained with an isotype control antibody and the resulting percentage of positive beads is shown. C) The limit of detection (LOD) for the PDAC^EV^ antibodies as single markers and an antibody cocktail is shown in both the minimum number and ng amount of EVs needed for BEAD flow. D) 1617 PDAC EVs were isolated on different days and were stained with an isotype control (orange) or EGFR (blue) antibody for BEAD flow analysis on different days to assess assay reproducibility.

Next, we used BEAD flow to test our previously identified PDAC^EV^ biomarker signature, a combination of EpCAM, EGFR, MUC1, GPC1, WNT-2^[10]^ on EVs collected from patient derived xenograft PDAC cell lines. We confirmed that each antibody comprising the signature showed high expression alone and in combination for flow cytometry detection (**Figure 3**). Importantly, in the absence of EVs, none of the antibodies used in these studies showed significant binding/signal on the streptavidin-PS beads when measured by flow cytometry (**Figure S3**). Using EVs derived from PDAC cell lines, each of the antibodies showed excellent positive signal by flow cytometry alone (**Figure 3A-D**)and in combination (**Figure 3F**), with the exception of GPC1 (**Figure 3E**). We have demonstrated GPC1 antibody signal in flow cytometry using biotinylated purified GPC1 protein attached to streptavidin PS beads (**Figure S4**). Low to moderate GPC1 expression was also observed in cell lines (**Figure 4A**), but little to no expression was seen on EV surfaces (**Figure 4B**). To ensure the functionality of the GPC1 antibodies, we tested antibodies from two different vendors with purified GPC1 protein as well as EVs from the Capan-2 cell line (**Figure S4**). While we observed high signal (> 1000 fold increase over isotype) with purified GPC1 proteins, little or no signal (< 1 fold) was observed in EVs, possibly because expression is below the detection limit of this assay.

**Figure 3.**
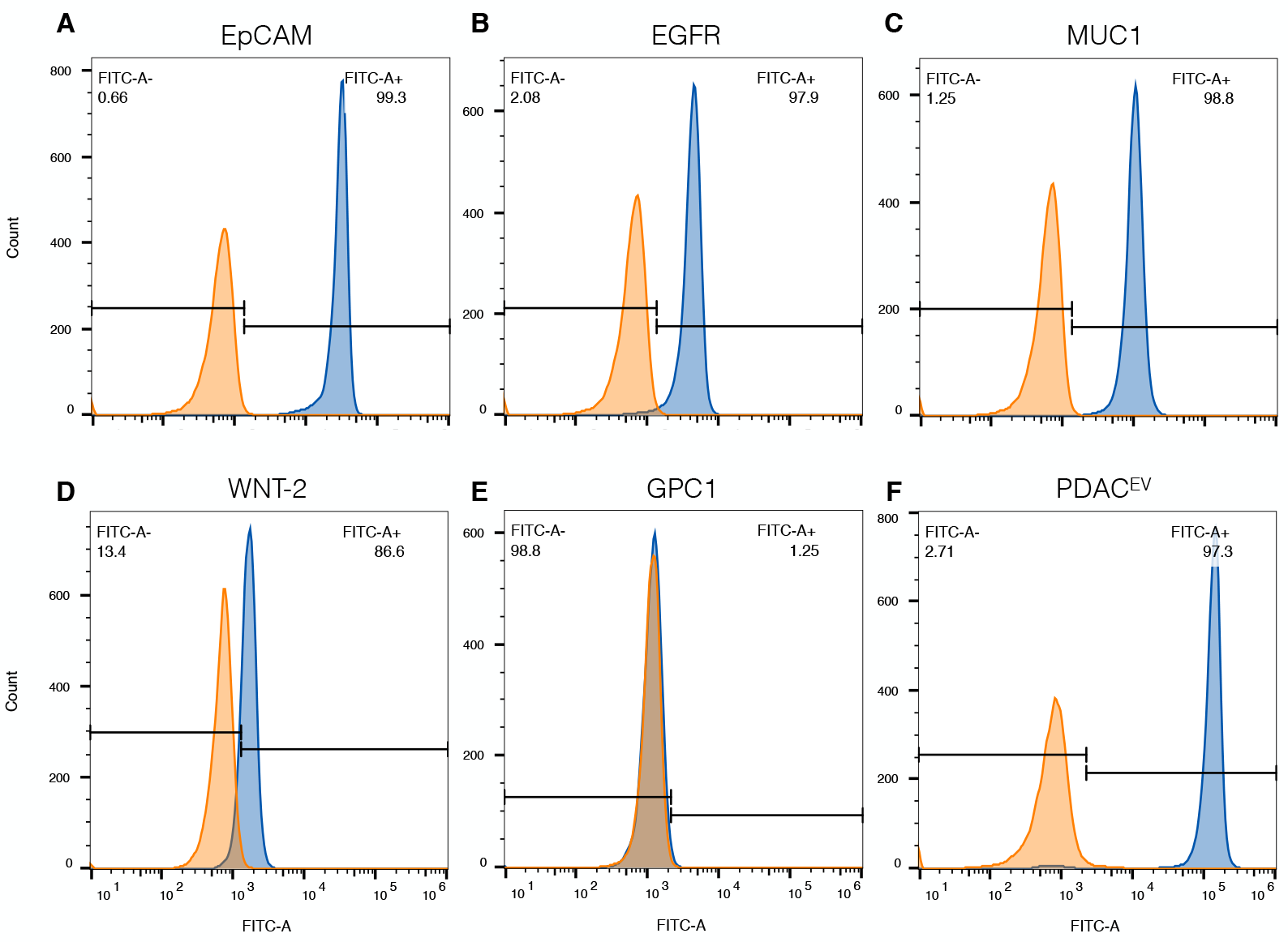
Validation of antibodies for BEAD flow. Biotinylated EVs were captured on 5 µm streptavidin polystyrene beads and stained with isotype control (orange) or primary antibodies (blue), followed by AlexaFluor 488 secondary antibodies. A) EpCAM, B) EGFR, C) MUC1, D) WNT-2, E) GPC1, F) PDAC^EV^ antibody cocktail (mixture of EpCAM, EGFR, MUC1, WNT-2, GPC1).

**Figure 4.**
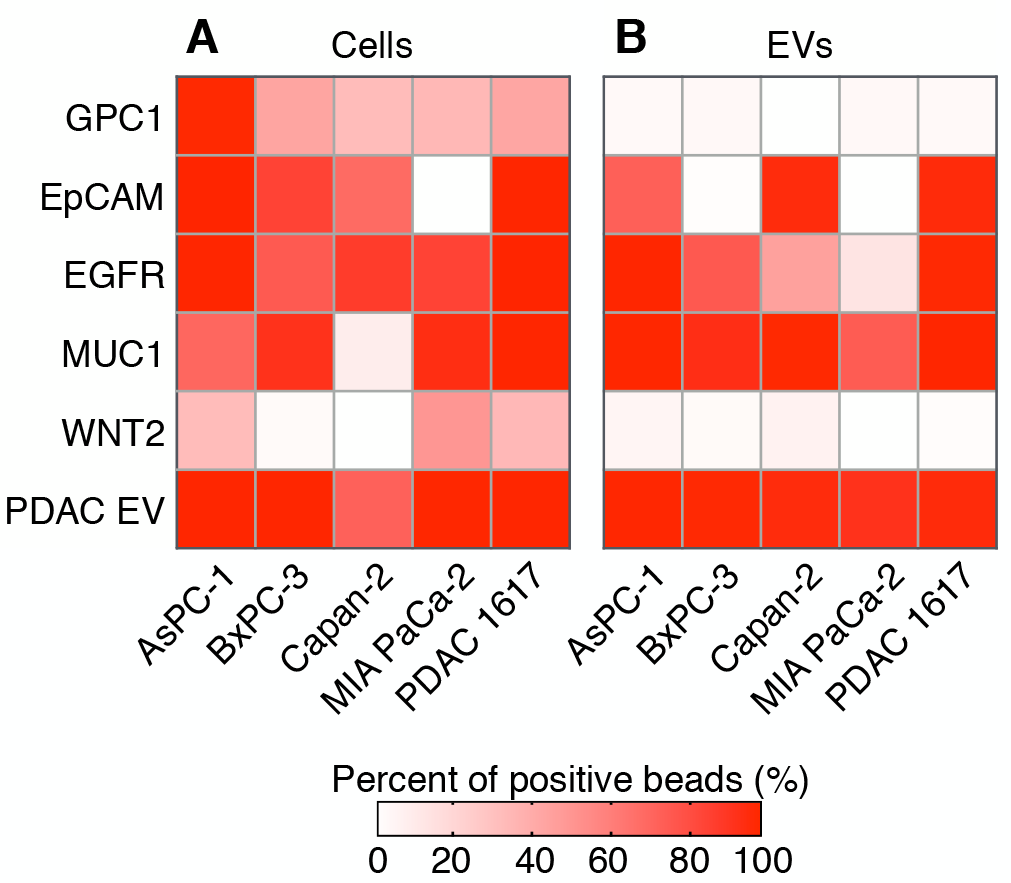
PDAC^EV^ biomarker expression in cell lines and EVs measured by BEAD flow. A) Cells were fixed and stained for surface biomarker expression using primary and AlexaFluor 488 secondary antibodies. B) EVs isolated from the same cell lines were biotinylated, captured on 5 µm streptavidin polystyrene beads and stained using the same antibodies as in A.

Comparing protein expression between EVs and their parental cells, we observed a moderate correlation (Pearson correlation coefficient r = 0.63; **Figure 4**) for five PDAC cell lines, including one patient-derived xenograft cell line (PDAC 1617). For outliers, such as EpCAM on BxPC-3 EVs, single marker expression levels could have fallen below our method’s limit of detection. Our PDAC^EV^ signature^[10]^ showed high signal for both EVs and their 5 parental cell lines.

Ultimately, the goal of BEAD flow is to analyze EVs from clinical samples. As such, EVs were isolated from plasma in a cohort of ten PDAC patients and three controls (**Figure 5**). Using our high throughput flow method, EV expression of eleven biomarkers were analyzed while consuming <1 mL plasma. Our previous five marker PDAC^EV^ signature showed good expression in PDAC EVs, with low to no expression in control EVs (**Figure 5**). The expression of five additional markers for pancreatic cancer based on recent literature reports (CD73,^[25]^ TIMP1,^[26]^ EphA2,^[19]^ LRG1,^[26]^ and Mesothelin^[27,28]^) were also analyzed, but none of the new markers surpassed the diagnostic accuracy of the PDAC^EV^ signature.^[10]^

**Figure 5.**
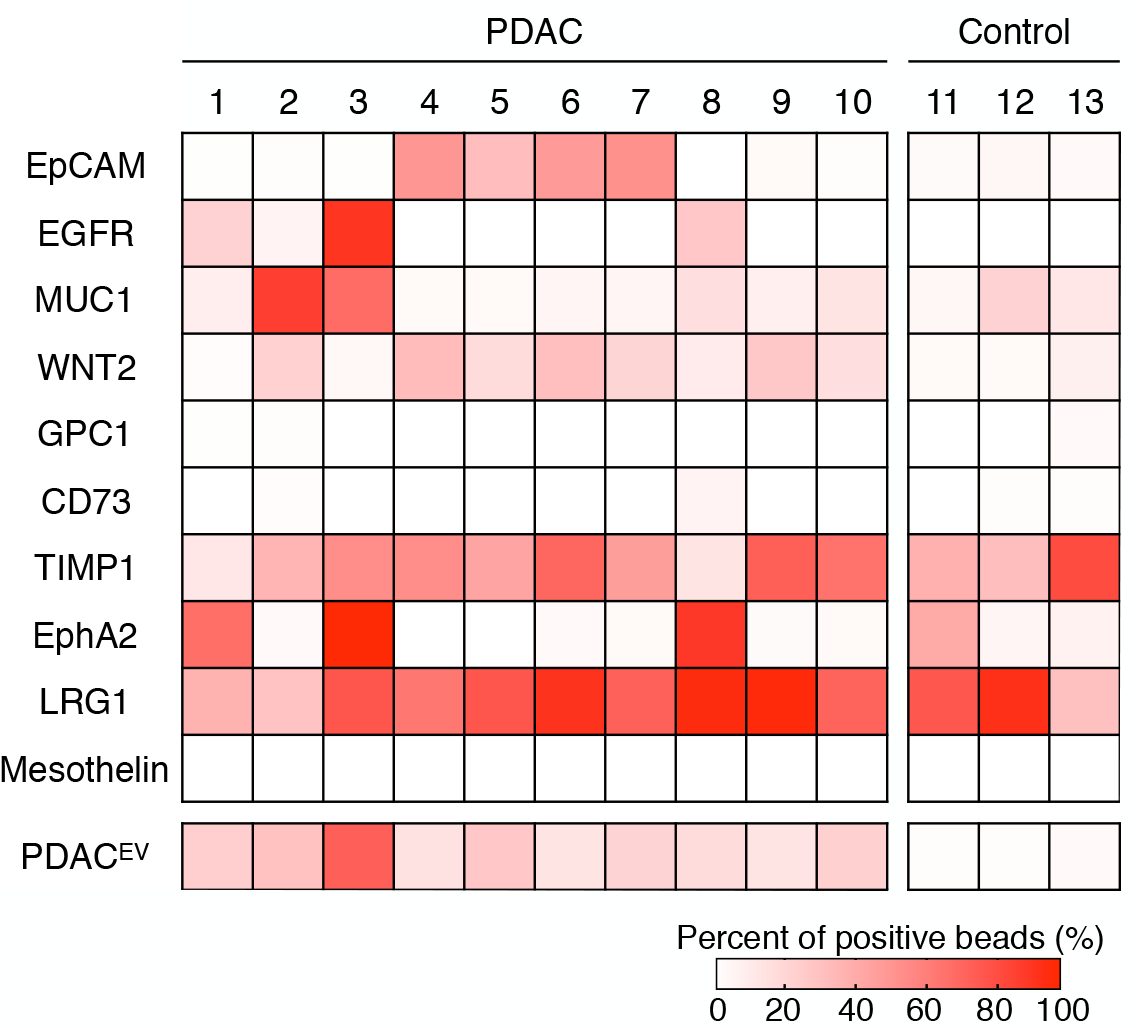
BEAD flow distinguishes PDAC from non-cancer patients. EVs were isolated from patient plasma using qEV columns (Izon), biotinylated, and captured on 5 µm streptavidin polystyrene beads. Beads were stained with the indicated primary antibodies (top heatmap)or with a PDAC^EV^ antibody cocktail (mixture of EpCAM, EGFR, MUC1, WNT-2, and GPC1; bottom heatmap). The percentage of positive beads is depicted in the heatmaps with P1-10 representing pancreatic cancer EVs and C1-3 representing controls.

For samples with very limited volumes, we tested whether an antibody cocktail of the PDAC^EV^ signature was useful instead of a linear combination of five individual markers. Using this antibody cocktail, we were able to readily distinguish PDAC from control samples (**Figure 5**). This strategy should allow us to analyze EVs from very small volumes of plasma (~100 μl) with high sensitivity and specificity for cancer versus non-cancerous states.

We have shown that integrating biotinylated EVs and streptavidin-coated beads results in enhanced sensitivity in a simplified assay format for analysis of EVs. Using BEAD flow, EVs from thirteen patient samples were analyzed for a PDAC^EV^ biomarker signature with excellent sensitivity. This method enables analysis from a simple patient blood draw and may be useful in identifying patients for further clinical assessment. While we focused on PDAC to develop this method, it is conceivable that it can be expanded to include additional biomarkers and cancer types. Combining bead enhanced EV analysis with minimal sample consumption aligns with scientific workflow needs for both clinical evaluations and research (e.g. biorepositories where precious samples are intended to be leveraged to their fullest potential).

## Experimental Section

### Cell Culture

AsPC-1, BxPC-3, Capan-2, and MIA PaCa-2 cells were obtained from ATCC. AsPC-1 and BxPC-3 cells were cultured in RPMI 1640 media, Capan-2 cells in McCoy’s 5a media, and MIA PaCa-2 cells in Dulbecco’s Modified Eagle’s medium (ThermoFisher). All media was supplemented with penicillin (10,000 IU)/streptomycin (10,000 μg/ml) (Corning Life Sciences) and 10% fetal bovine serum (Atlanta Biologics). 1617 cells are a patient-derived xenograft cell line (kind gift from Dr. Carlos Fernandez del Castillo, Massachusetts General Hospital) and were cultured in a 50:50 mix of DMEM/F-12 media, supplemented as above.

### EV Isolation from Cell Culture

Cells were trypsinized and split into 8-10, 15-cm dishes in appropriate media containing 5% exosome-depleted FBS (ThermoFisher) and grown for 48-72 hours. Media containing released EVs was collected and centrifuged at 300 × g for 10 min, followed by filtration through a 0.22 µm cellulose-acetate vacuum filter. Conditioned media then underwent ultracentrifugation at 100,000 × g for 70 min (4°C). Supernatant was removed, EV pellets resuspended in a single tube in PBS and centrifuged a second time at 100,000 × g for 70 min. Supernatant was again removed and the EV pellet resuspended in ~100 µl PBS. EVs were stored at -80°C until use.

### EV Isolation from Plasma

Plasma was collected from patients who provided informed written consent under a sample collection protocol approved by the Dana-Farber/Harvard Cancer Center IRB. PDAC patients had locally advanced disease, with samples collected at baseline prior to initiation of systemic treatment. Whole blood was collected in one 10 mL purple-top EDTA tube, mixed by inverting 10 times, and then centrifuged for 10 min at 400 × g (4°C). The plasma layer was collected in a 15 mL conical tube without disturbing the buffy coat and was then centrifuged for 10 min at 1100 × g (4°C). The plasma layer was pipetted into a 15 mL tube and stored at - 80°C until processing for EVs. qEV size-exclusion columns (Izon) were used for EV isolation and were first washed with 10 mL PBS. During column washing, 500 μl plasma was cleared by centrifugation at 1500 × g for 10 min (4°C). The supernatant was then spun again at 10,000 × g for 20 min (4°C). Cleared supernatant was loaded onto the qEV column and 0.5 mL fractions were immediately collected. As soon as the sample completely entered the resin, PBS was added. The first six fractions were discarded (3 mL, column void volume). Fractions 7-9 (1.5 mL total) were pooled and filtered through an Ultrafree 0.22 µm centrifugal filter at 12,000 × g for 1 min. The pooled fractions were then concentrated using an Amicon Ultra-4 10kDa filter by centrifugation at 3200 × g for 15 min. Concentrated EVs were stored at -80°C.

### EV Biotinylation

Total EV protein was measured using 5-10 μl EVs and the Qubit protein assay kit (ThermoFisher). Samples were biotinylated for 30 min with a 20-fold molar excess of EZ-Link NHS-PEG4-Biotin (ThermoFisher) in 100-200 μl total volume. Excess biotin was removed using MW 3000 Exosome Spin Columns (ThermoFisher). Total biotinylated EV protein was again measured with the Qubit protein assay.

### EV Capture on Beads

10 μl of 5.0-5.9 μm streptavidin coated polystyrene particles (Spherotech) were diluted in 1 mL PBS/1% BSA, centrifuged at 3000 × g for 2 min and supernatant was removed. Beads were mixed in a total of 10 µl with 500 ng EVs and PBS. For more than one sample of the same EV population, EVs were captured on beads in batch (ex: 5 µg EVs in 100 µl PBS/beads) and later split into multiple tubes for antibody staining. EVs were incubated for 30 min with beads on a HulaMixer (ThermoFisher). Excess reacted EVs were removed by washing twice with PBS/1% BSA. EV-beads were diluted in 100 µl PBS/1% BSA and transferred to a 96- well u-bottom plate for antibody staining.

### Single EV Flow Cytometry Staining and Analysis

2 µg EVs were incubated with 1 µl isotype control or EGFR-FITC antibodies overnight with rotation/mixing. Excess antibody was removed by purifying EVs using the qEV size exclusion columns (Izon), as described above for EV purification from patient plasma. Samples were run on the CytoFlex flow cytometer using Violet SSC to resolve smaller particles and the following settings: SSC-A 30, VSSC-A 22 (threshold 2000 on area), FITC-A 20, FSC 201.

### Flow Cytometry Staining and Analysis

EVs and cells were stained using the same antibodies and procedure. Cells were prepared for flow staining by fixing 500,000 cells per antibody condition in 4% formaldehyde (ThermoFisher) in PBS for 15 min at room temp on a nutating mixer. Cells were washed in PBS/1% BSA and aliquoted to a 96-well u-bottom plate for staining. Cells or EVs were pelleted in the 96-well plate and excess buffer was removed by flicking over a sink. Samples were resuspended in PBS/1% BSA or in 10 µg/ml primary antibody diluted in the same buffer and incubated for 30 min on a plate shaker set to medium speed (see **Table S2** for a complete list of antibodies used in this study). Cells or EVs were pelleted and washed twice with 150 µl PBS/1% BSA. Cells or EVs were then resuspended in the appropriate AlexaFluor 488 secondary antibody diluted 1:1000 in PBS/1% BSA and incubated for 30 min (protected from light) on a plate shaker. Samples were again pelleted, washed twice with 150 µl PBS/1% BSA, and resuspended in 200 µl PBS/1% BSA for flow analysis. Samples were measured using a Beckman Coulter CytoFlex flow cytometer with 96-well plate handling using the following settings: cells FSC 49V, SSC 104V, FITC 20V; EVs FSC 201V, SSC 90V, FITC 159V. Sample backwash and mixing were turned off and a total of 10,000 events were collected. Samples were analyzed by gating on the isotype control population and measuring the percentage of positive events greater than the isotype control.

### Intensity per Bead Calculation

The average bead intensity can be calculated by the difference between the primary antibody treated beads and isotype controls through the integration of intensity area divided by counted bead number (**Figure S5**). The bead intensity used herein can be expressed as: 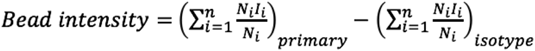

All data is processed through FlowJo and Matlab.

## Supporting Information

Supporting Information is available at the end of the document.

## Acknowledgements

Hsing-Ying Lin and Katherine S. Yang contributed equally to this work. The authors acknowledge funding from the U.S. National Institutes of Health (NIH) grants NIH R01CA204019 (R.W.), P01CA069246 (R.W.), 4R00CA201248 (H.I.), R01HL113156 (H.L.), R21CA205322 (H.L.), U01 CA210171 (B.M.W.), a grant from the Lustgarten Foundation (R.W.), a grant from the Hale Center for Pancreatic Cancer Research (B.M.W.) and a pilot grant from the Andrew L. Warshaw, M.D. Institute for Pancreatic Cancer Research at MGH (K.S.Y. and H.I.).

## Supporting Information

### Bead Enhancement of EV Analysis

**Figure S1.**
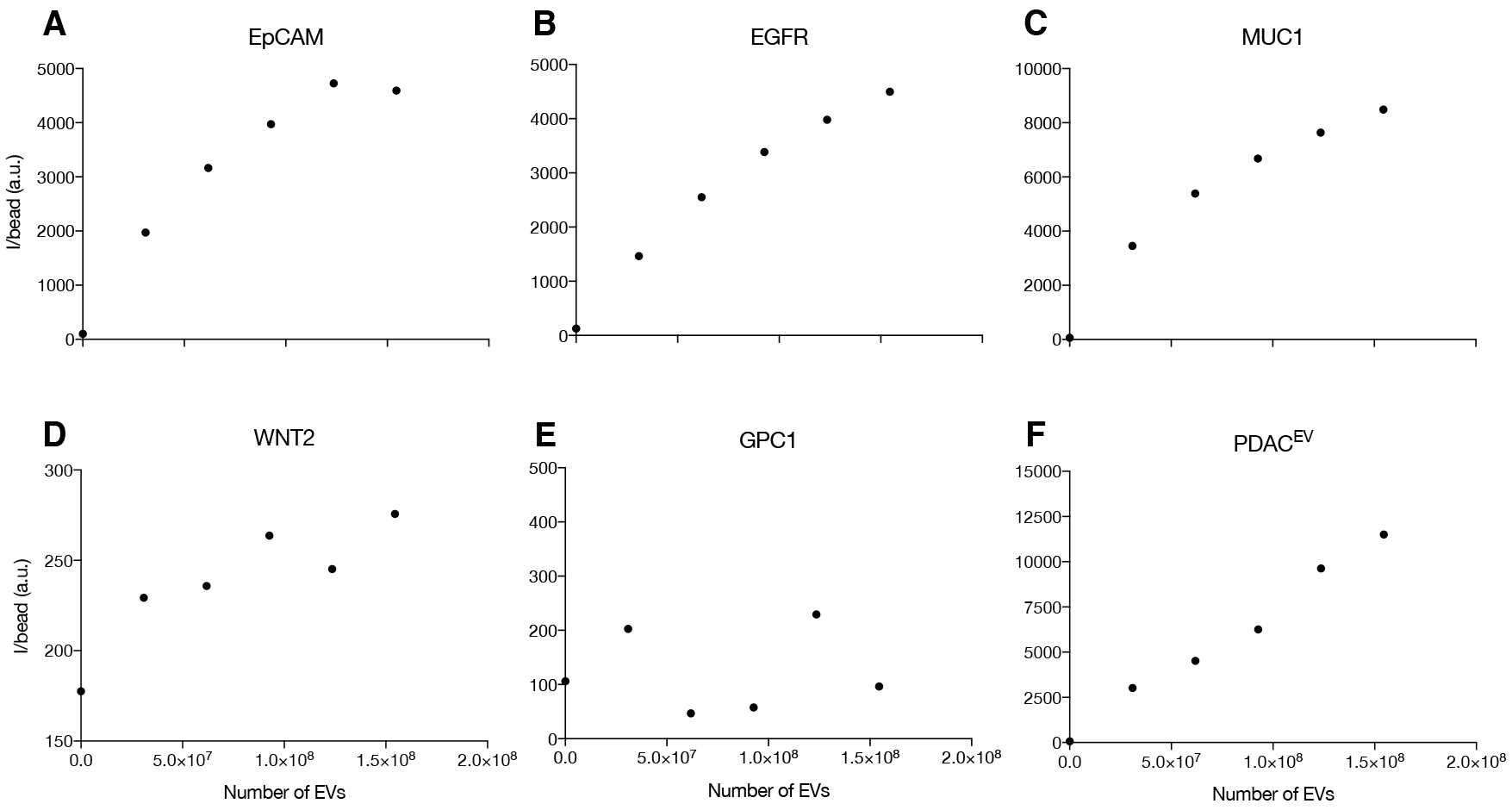
EV titration curves for limit of detection analysis of each antibody used in BEAD flow. Increasing numbersof biotinylated 1617 EVs were captured on streptavidin polystyrene beads and stained with antibodies against (A) EpCAM, (B) EGFR, (C) MUC1, (D) WNT2, (E) GPC1, and (F) PDAC^EV^ cocktail (mixture of all 5 antibodies). Signal intensity per bead was calculated, with isotype control signal subtracted, and plotted versus the number of EVs.

**Table S1.** Minimum EV amounts needed from 1617 PDAC EVs in BEAD flow for antibody detection.

| PDAC<sup>EV</sup> Antibody | Limit of Detection (ng)<sup>a)</sup> | Limit of Detection (# EVs) |
| --- | --- | --- |
| Cocktail | 74.24 | 2.29 x 10<sup>7</sup> |
| EGFR | 41.11 | 1.27 x 10<sup>7</sup> |
| EpCAM | 86.56 | 2.67 x 10<sup>7</sup> |
| MUC1 | 86.61 | 2.68 x 10<sup>7</sup> |
| WNT-2 | 127.45 | 3.94 x 10<sup>7</sup> |
| GPC1<sup>b)</sup> | 15042.39 | 4.65 x 10<sup>9</sup> |

<sup>a)</sup>ng: the total protein amount of biotinylated EVs required as measured using the Qubit protein assay; <sup>b)</sup>GPC1 expression is below the limit of detection on 1617 PDAC EVs.

**Figure S2.**
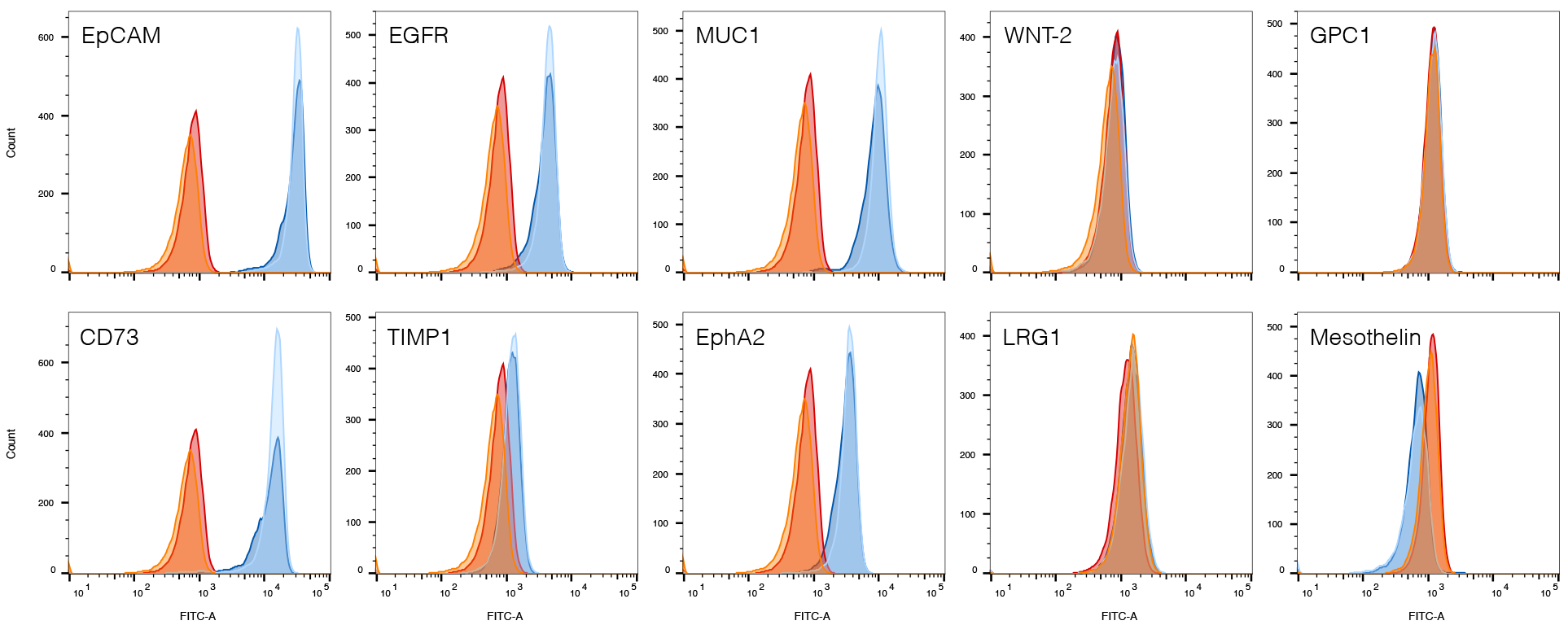
BEAD flow reproducibility. 1617 PDAC EVs were isolated and biotinylated on different days. EVs were captured on streptavidin polystyrene beads and were stained with an isotype control (orange) or primary (blue) antibody for BEAD flow analysis on different days to assess assay reproducibility.

**Figure S3.**
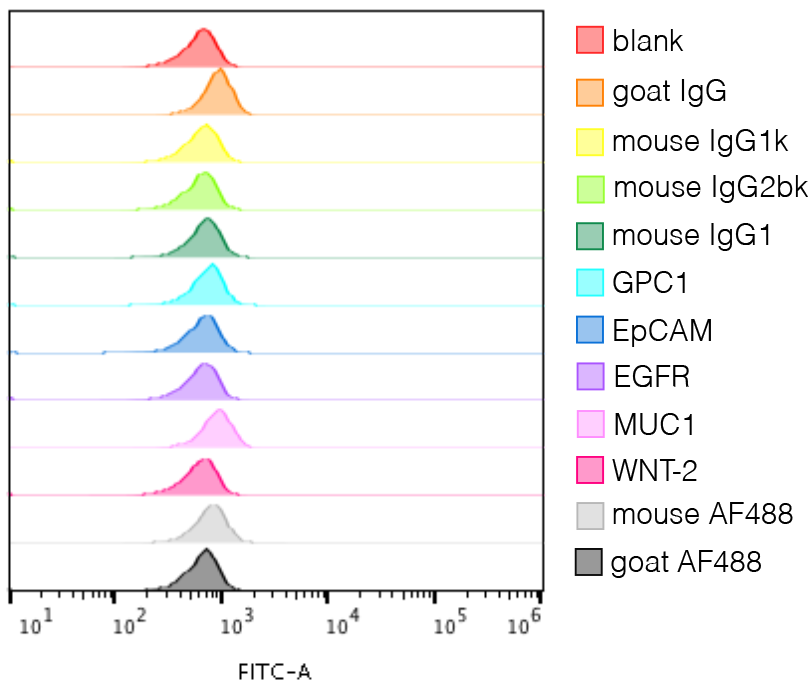
Background antibody binding to streptavidin polystyrene beads in BEAD flow. 5 µm streptavidin polystyrene beads were stained with the indicated primary antibodies, followed by the indicated secondary antibodies to confirm that the antibodies used in this study do not bind to beads in the absence of EVs.

**Figure S4.**
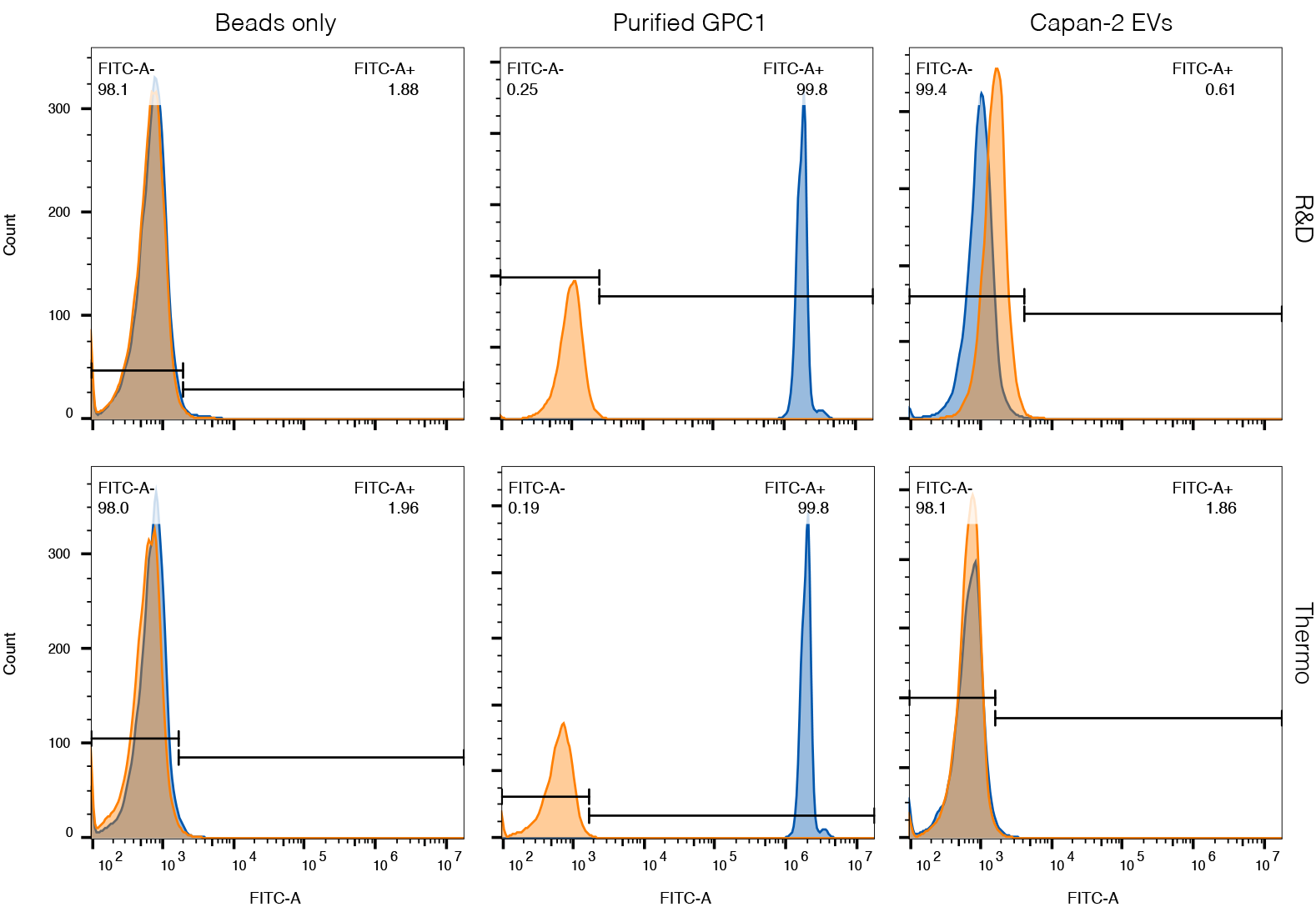
GPC1 antibody testing in BEAD flow. Streptavidin polystyrene beads alone (left), with biotinylated purified GPC1 protein (middle), or with biotinylated Capan-2 EVs (right) captured were stained with two commercially available GPC1 antibodies (blue) and compared to matching isotype control antibodies (orange).

**Figure S5.**
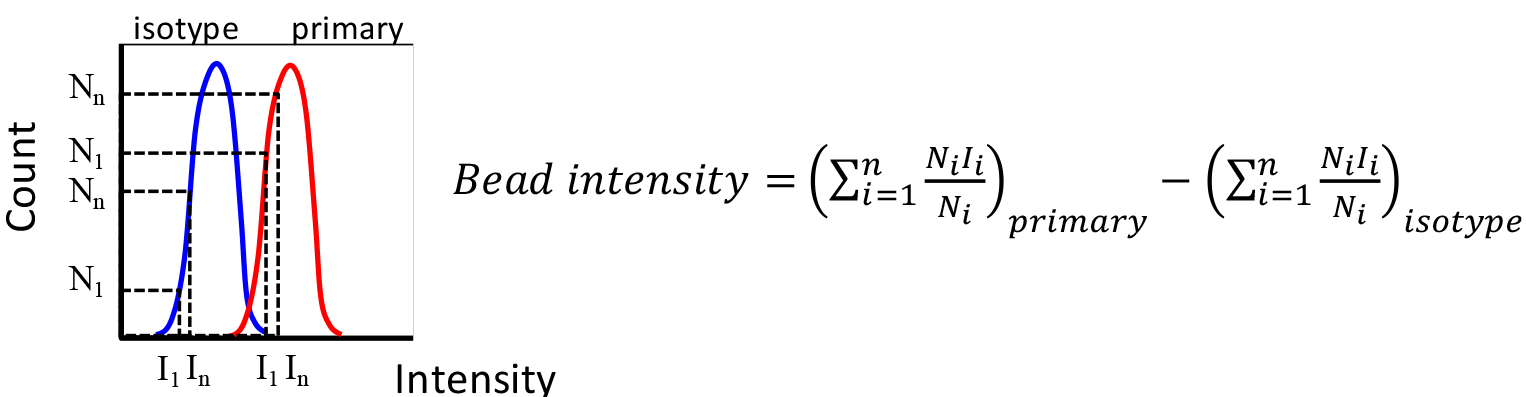
Average bead intensity calculation. The average bead intensity is calculated by the difference between theprimary antibody treated beads and isotype control beads through the integration of intensity area divided by counted bead number.

**Table S2.** Antibodies used in this study.

|  | Antibody | Company | Catalog # | Species | Clonality | Use |
| --- | --- | --- | --- | --- | --- | --- |
| Isotype | Goat IgG | Invitrogen | 02-6202 | Goat |  |  |
|  | Mouse IgG1k | BioLegend | 400102 | Mouse |  | FC, WB, IP, ICC, IF, IHC |
|  | Mouse IgG2bk | Abcam | Ab18469 | Mouse | Monoclonal | IHC, WB, FC, IP |
|  | Mouse IgG1 | R&D | MAB002 | Mouse | Monoclonal | FC |
|  | Rabbit IgG | Abcam | Ab172730 | Rabbit | Monoclonal | IF, IHC, FC, CHIPseq, IP |
|  | Rat IgG2ak | BioLegend | 400502 | Rat |  | FC, IF, IHC, WB, IP |
|  | Rat IgG2ak-AF488 | BioLegend | 400525 | Rat |  | FC |
| Primary | Glypican-1 (GPC1) | R&D | AF4519 | Goat IgG | Polyclonal | WB, FC, IF |
|  | EpCAM | Abcam | Ab20160 | Mouse IgG1k | Monoclonal | IF, FC, ELISA, IHC |
|  | EGFR | Abcam | Ab30 | Mouse IgG2bk | Monoclonal | IHC, IP, IF, FC |
|  | MUC1 | Fitzgerald | 10-M93A | Mouse IgG1 | Monoclonal | ELISA, IHC, WB |
|  | WNT-2 | Santa Cruz | Sc-514382 | Mouse IgG1k | Monoclonal | WB, IP, IF, ELISA |
|  | CD73 | BD | 550256 | Mouse IgG1k | Monoclonal | FC |
|  | EphA2 | R&D | MAB3035 | Mouse IgG<sub>2A</sub> | Monoclonal | WB, FC, IF |
|  | Mesothelin | R&D | MAB32652 | Rat IgG<sub>2A</sub> | Monoclonal | FC |
|  | LRG1 | Invitrogen | PA5-25904 | Rabbit IgG | Polyclonal | WB, IHC, FC |
|  | TIMP1 | Invitrogen | MS608PABX | Mouse IgG1 | Monoclonal | FC, IF, IHC, WB |
|  | EGFR-FITC | Abcam | Ab11400 | Rat IgG<sub>2A</sub> | Monoclonal | FC |
| Secondary<br>AlexaFluor 488 | Mouse IgG (H+L) | Abcam | Ab150117 | Mouse IgG | Polyclonal | IHC, IF, FC, ELISA |
|  | Goat IgG (H+L) | Abcam | Ab150133 | Goat IgG | Polyclonal | IHC, IF, FC, ELISA |
|  | Rabbit IgG (H+L) | Abcam | Ab150073 | Rabbit IgG | Polyclonal | IF, IC, IHC, ELISA |
|  | Rat IgG (H+L) | Abcam | Ab150153 | Rat IgG | Polyclonal | IF, ELISA, FC, IHC |

## References

[1] A. M. Lennon, C. L. Wolfgang, M. I. Canto, A. P. Klein, J. M. Herman, M. Goggins, E. K. Fishman, I. Kamel, M. J. Weiss, L. A. Diaz, N. Papadopoulos, K. W. Kinzler, B. Vogelstein, R. H. Hruban, Cancer Res. 2014, 74, 3381.

[2] S. Kaur, M. J. Baine, M. Jain, A. R. Sasson, S. K. Batra, Biomark Med. 2012, 6, 597.

[3] E. L. Fogel, S. Shahda, K. Sandrasegaran, J. DeWitt, J. J. Easler, D. M. Agarwal, M. Eagleson, N. J. Zyromski, M. G. House, S. Ellsworth, I. El Hajj, B. H. O’Neil, A. Nakeeb, S. Sherman, Am. J. Gastroenterol. 2017, 112, 537.

[4] S. K. Maithel, S. Maloney, C. Winston, M. Gönen, M. I. D’Angelica, R. P. Dematteo, W.R. Jarnagin, M. F. Brennan, P. J. Allen, Ann. Surg. Oncol. 2008, 15, 3512.

[5] J. Martinez-Useros, J. Garcia-Foncillas, Biomed. Res. Int.. 2016, 2016, 4873089.

[6] J. E. Kim, K. T. Lee, J. K. Lee, S. W. Paik, J. C. Rhee, K. W. Choi, J. Gastroenterol. Hepatol. 2004, 19, 182.

[7] S. Kawai, K. Suzuki, K. Nishio, Y. Ishida, R. Okada, Y. Goto, M. Naito, K. Wakai, Y. Ito, N. Hamajima, Int. J. Cancer. 2008, 123, 2880.

[8] M. Herreros-Villanueva, L. Bujanda, Ann. Transl. Med. 2016, 4, 134.

[9] S. A. Melo, L. B. Luecke, C. Kahlert, A. F. Fernandez, S. T. Gammon, J. Kaye, V. S. LeBleu, E. A. Mittendorf, J. Weitz, N. RahbariC. Reissfelder, C. Pilarsky, M. F. Fraga, D. Piwnica-Worms, R. Kalluri, Nature. 2015, 523, 177.

[10] K. S. Yang, H. Im, S. Hong, I. Pergolini, A. F. Del Castillo, R. Wang, S. Clardy, C. H. Huang, C. Pille, S. Ferrone, R. Yang, C. M. Castro, H. Lee, C. F. Del Castillo, R. Weissleder, Sci. Transl. Med. 2017, 9, eaaI3226.

[11] C. Théry, L. Zitvogel, S. Amigorena, Nat. Rev. Immunol. 2002, 2, 569.

[12] G. Camussi, M. C. Deregibus, S. Bruno, C. Grange, V. Fonsato, C. Tetta, Am. J. Cancer Res. 2011, 1, 98.

[13] J. Rak, A. Guha, Bioessays. 2012, 34, 489.

[14] H. Shao, J. Chung, L. Balaj, A. Charest, D. D. Bigner, B. S. Carter, F. H. Hochberg, X. O. Breakefield, R. Weissleder, H. Lee, Nat. Med. 2012, 18, 1835.

[15] H. Shao, J. Chung, K. Lee, L. Balaj, C. Min, B. S. Carter, F. H. Hochberg, X. O. Breakefield, H. Lee, R. Weissleder, Nat. Commun. 2015, 6, 6999.

[16] H. Im, H. Shao, Y. I. Park, V. M. Peterson, C. M. Castro, R. Weissleder, H. Lee, Nat. Biotechnol. 2014, 32, 490.

[17] K. R. Jakobsen, B. S. Paulsen, R. Bæk, K. Varming, B. S. Sorensen, M. M. Jørgensen, J. Extracell. Vesicles. 2015, 4, 26659.

[18] B. Madhavan, S. Yue, U. Galli, S. Rana, W. Gross, M. Müller, N. A. Giese, H. Kalthoff, T. Becker, M. W. Büchler, M. Zöller, Int. J. Cancer. 2015, 136, 2616.

[19] K. Liang, F. Liu, J. Fan, D. Sun, C. Liu, C. J. Lyon, D. W. Bernard, Y. Li, K. Yokoi, M. H. KatzE. J. Koay, Z. Zhao,Y. Hu, Nat. Biomed. Eng. 2017, 1, 0021.

[20] G. Kibria, E. K. Ramos, K. E. Lee, S. Bedoyan, S. Huang, R. Samaeekia, J. J. Athman, C.V. Harding, J. Lötvall, L. Harris, C. L. Thompson, H. Liu, Sci. Rep.. 2016, 6, 36502.

[21] A. Morales-Kastresana, B. Telford, T. A. Musich, K. McKinnon, C. Clayborne, Z. Braig, A. Rosner, T. Demberg, D.C. Watson, T. S. Karpova, G. J. Freeman, R. H. DeKruyff, G.N. Pavlakis, M. Terabe, M. Robert-Guroff, J. A. Berzofsky, J. C. Jones, Sci. Rep. 2017, 7, 1878.

[22] E. van der Pol, F. A. Coumans, A. E. Grootemaat, C. Gardiner, I. L. Sargent, P. Harrison, A. Sturk, T. G. van Leeuwen, R. Nieuwland, J. Thromb. Haemost. 2014, 12, 1182.

[23] A. Morales-Kastresana, J. C. Jones, Methods Mol. Biol. 2017, 1545, 215.

[24] C. Théry, S. Amigorena, G. Raposo, A. Clayton, Curr. Protoc. Cell Biol. 2006, Chapter 3, Unit 3.22.

[25] B. Zhang, Cancer Res. 2010, 70, 6407.

[26] M. Capello, L. E. Bantis, G. Scelo, Y. Zhao, P. Li, D.S. Dhillon, N.J. Patel, D.L. Kundnani, H. Wang, J. L. Abbruzzese, A. Maitra, M.A. Tempero, R. Brand, M.A. Firpo, S.J. Mulvihill, M.H. Katz, P. Brennan, Z. Feng, A. Taguchi, S.M. Hanash, J. Natl. Cancer Inst. 2017, 109,djw266.

[27] P. Argani, C. Iacobuzio-Donahue, B. Ryu, C. Rosty, M. Goggins, R.E. Wilentz, S.R. Murugesan, S.D. Leach, E. Jaffee, C.J. Yeo, J.L. Cameron, S.E. Kern, R.H. Hruban, Clin. Cancer Res. 2001, 7, 3862.

[28] M. Li, U. Bharadwaj, R. Zhang, S. Zhang, H. Mu, W.E. Fisher, F.C. Brunicardi, C. Chen, Mol. Cancer Ther. 2001, 7, 286.

